# CoalMiner: a coalescent model generator for *fastsimcoal2*

**DOI:** 10.64898/2026.06.25.734618

**Authors:** Raya Esplin-Stout, Arun Sethuraman

## Abstract

Demographic inference using the Site Frequency Spectrum (SFS) is often constrained by the number and complexity of models tested. Here we present a coalescent model generator called *CoalMiner* for use with *fastsimcoal2*. *CoalMiner* utilizes a decision tree framework to generate biologically plausible models, with user input dictating the number and ranges of demographic parameters and histories, which can then be plugged into the *fastsimcoal2* pipeline. Using extensive simulations and empirical data, we show that *CoalMiner* is an effective helper tool to explore demographic model space. *CoalMiner* is written in Python and is freely available on GitHub: https://github.com/raywray/coalminer with numerous vignettes and tutorials.

## Introduction

Methods used to estimate the demographic history of species and populations have evolved with the growth of modern sequencing technology. From single-locus haplotype-based inference (Beerli & Felsenstein, 2001; Chung & Hey, 2017; Hey et al., 2018) to inference from millions of variants (Excoffier & Foll, 2011; Gutenkunst et al., 2009), these methods are utilized extensively across a variety of biological systems and applications (Excoffier et al., 2021; Gutenkunst, 2021; Tran et al., 2024). Specifically, methods using the site frequency spectrum (SFS) for evolutionary inference have become increasingly popular within the last decade, due to the shapes of the SFS being sensitive to demographic processes and signatures of selection across timescales (Smith & Hahn, 2024), the ease with which demographic and selection parameters can be inferred using statistical approaches applied to the SFS (Keightley & Halligan, 2011), and the simplicity of representing population-scale genomic data from approaches like Restriction-side Associated DNA sequencing (RADseq) and Single Nucleoptide Polymorphism (SNP) arrays as multidimensional histograms of allele frequencies (de Jong et al., 2021).

Methods using the SFS are computationally efficient, accurate, and have broad applications in a variety of biological fields, especially when combined with model-based inference of evolutionary histories using recent tools such as *fastsimcoal2* (Excoffier et al., 2021) and *dadi* (Gutenkunst et al., 2009). For instance, *fastsimcoal2* has been used to identify prehistoric population expansions and domestication signatures in yak (Qiu et al., 2015), estimate population fragmentation in endangered Iberian grasshoppers (González-Serna et al., 2018), reconstruct recent takin divergence histories for conservation planning (Yang et al., 2022), and to infer complex demographic histories among European whitefish populations (Crotti et al., 2021) in fisheries research. In agriculture, *fastsimcoal2* was used to reveal population splits, migration, and bottlenecks in *Coffea arabica* (Salojärvi et al., 2024), and to resolve the evolutionary relationships of wild yams, correcting a previously published phylogenetic finding (Sugihara et al., 2020). In human evolution, *fastsimcoal2* served as the simulation engine within an ABC-DL framework used to model North African demographic history, supporting the inference that Arab and Amazigh populations have distinct evolutionary origins (Serradell et al., 2024).

However, users of SFS-based evolutionary inference tools are limited in the number of models they can feasibly test, since each model requires explicit specification of topology, parameter boundaries, and priors. Contemporary population structure and diversity, as inferred from the SFS, are also molded by numerous factors such as ancestral population sizes, divergences, and migration (gene flow) from extant and extinct species and populations, over thousands, and even millions of years (Noskova et al., 2020). As a result, most studies, including those listed above, only test a handful of evolutionary demographic models. Existing tools for automated model exploration, such as *GADMA* and *GADMA2* (Noskova et al., 2020, 2023), utilize genetic algorithm-based global searches over the demographic model space. However, these tools are built around the *dadi* and *moments* (Jouganous et al., 2017) inference frameworks and therefore do not produce input files compatible with *fastsimcoal2*. No existing tool automates the generation of *fastsimcoal2*-native demographic topologies, leaving users to hand-specify each .*tpl* and .*est* file. This manual process is time-consuming, error-prone, and practically limits the number of models evaluated. While prior knowledge of the biology of the species of interest is often utilized in designing these limited models, we hypothesize that broader exploration of demographic model space may improve the robustness and biological plausibility of demographic inference. Indeed, parametric models that are too simple may fail to capture key demographic events, leading to biased or misleading inferences (Adrion et al., 2020).

Here, we introduce a companion software, *CoalMiner*, that generates a multitude of possible evolutionary models from simple user input, which can be used as input for evolutionary inference using *fastsimcoal2*. We anticipate that *CoalMiner* will alleviate model-generation limitations and enable a more comprehensive exploration of demographic model space, leading to more accurate and robust evolutionary inference. *CoalMiner* is a stochastic demographic model generation framework for *fastsimcoal2*, and we evaluate its performance here using both simulated benchmark models and empirical population genomic data from domesticated hops (*Humulus lupulus L*.).

## Materials & Methods

### *CoalMiner* Model Generation Pipeline

*CoalMiner* assembles demographic models that may include a range of historical events, with a decision-tree-based algorithm. During topology generation, *CoalMiner* decides based on a “coin flip” whether to add the following events: ghost population, migration, admixture, and bottleneck events, and determines the order of historical events. Historical events are generated to ensure logical consistency (e.g., populations cannot reappear after merging) and are inserted in a time-consistent order. If selected, migration matrices are constructed to reflect model structure, with event-aware connectivity constraints.

Each model is written as a .*tpl* file, and parameter ranges are defined in a corresponding .*est* file. Simple and complex parameters are then assigned based on the .*tpl* file, user-specified distributions, and event structure. Figure 1 describes the basic workflow of *CoalMiner*.

**Figure 1:**
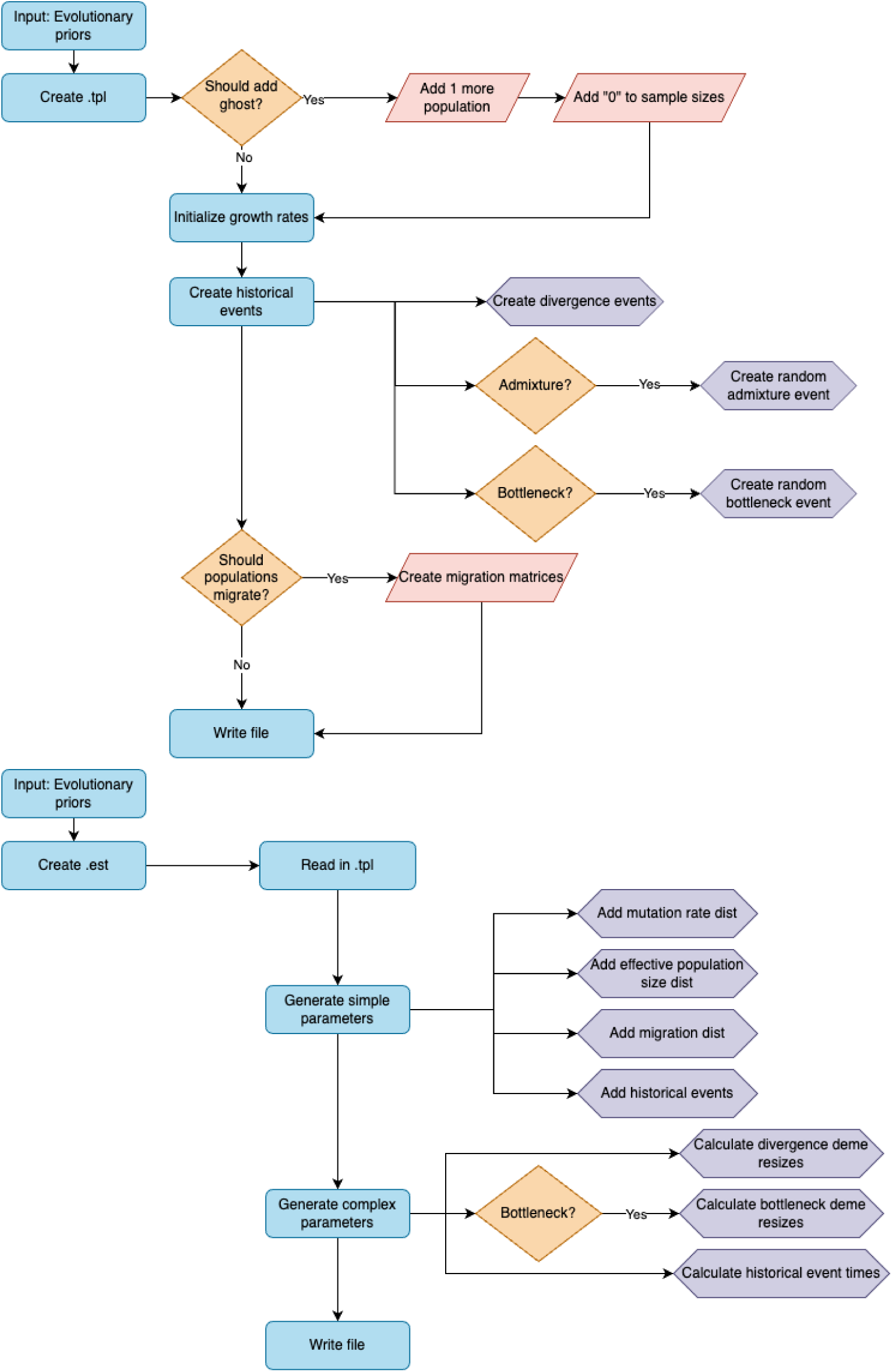
The decision tree describing the *CoalMiner* model generation pipeline for producing input files compatible with *fastsimcoal2*. The pipeline consists of two sequential modules. The top module generates a topology file (.*tpl*), which generates population topology and historical events. Beginning with user-specified evolutionary priors, *CoalMiner* first determines stochastically (50% probability each) whether to include a ghost population and, if selected, adds another population with a sample size of zero. Growth rates are initialized for all populations. Historical events are then generated either deterministically (divergence events are always created) or stochastically (admixture and bottleneck events are each included with a 50% probability). If migration is selected (50% probability), migration matrices are constructed with event-aware connectivity, completing the .*tpl*. The bottom module generates the matching .*est* file, which encodes parameter distributions. The generated .*tpl* file is read in, and simple parameters are assigned for mutation rate, effective population sizes, migration rates, and historical event times. Complex parameters are then computed, including divergence and bottleneck deme resizes (calculated as population-size ratios) and historical event times (calculated as cumulative sums of spacing intervals to enforce chronological order). Colors denote node type: blue rectangles indicate processes, orange diamonds indicate stochastic decisions, pink parallelograms indicate outputs or additions, and purple hexagons indicate parameter calculations.

### Detailed *CoalMiner* Algorithm

For each model, *CoalMiner* creates a .*tpl* file describing the population topology and events, and a matching .*est* file specifying parameter distributions. Both the number of models and parameter priors are defined in a user-provided YAML configuration file.

#### .tpl File Generation

For the population structure, the user specifies the number of observed populations. With 50% probability, a ghost (unsampled) population is added. Sample sizes are assigned to observed populations; ghost populations receive size “0.”

Historical events begin with divergence (or coalescence, backwards in time). All populations are first partitioned into “sources” and “sinks.” *CoalMiner* then iterates through the source-sink assignments, generating one divergence event per source. Each divergence event includes the source and sink population indices, a full lineage collapse (migrants = 1), a growth rate of 0, and a population resize that is either a RELANC parameter (50% probability) or 1, where a value of 1 indicates no resize.

Divergence events are generated in an order that ensures logical backward-time ancestry. This process continues until only one population remains (in other words, all populations have successfully coalesced into a single lineage).

With 50% probability, a single admixture event is added. Multiple candidate sources and sinks are randomly generated from all available populations, and a unique source-sink pair is selected from these candidates. A random admixture proportion (0 ≤ m ≤ 1) is assigned. The admixture event is inserted into the historical event list in a time-consistent position relative to existing divergence events.

With 50% probability, a single bottleneck is added. A population is randomly selected to undergo a temporary reduction in size. The onset event is generated first, including a RESBOT deme resize parameter. The corresponding recovery event (RESBOTEND) is then constructed automatically from the onset event during the ordering step and inserted immediately after it in the historical event list. Together, the two events bracket the bottleneck period.

All historical events (divergence, admixture, and bottleneck) are then passed to an ordering function that places admixture and bottleneck events into chronologically valid positions relative to the divergence scaffold. Temporal consistency is enforced during construction rather than by post-hoc filtering. As a note, there will be no events between a bottleneck start and end; those will be consecutive each time.

If migration is selected (50% probability), a migration matrix is generated. With a further 50% probability, migration rates vary across matrices: when variation is enabled, each matrix receives uniquely named parameters suffixed with the matrix index (e.g., MIGxy_1, MIGxy1, MIGxy_2, MIGxy2); when variation is disabled, all matrices share the same parameter names. In both cases, migration between coalesced populations is disabled by zeroing out the corresponding matrix entries following each divergence event.

#### .est File Generation

For simple parameters, mutation rate, effective population sizes, migration rates, and event times are sampled from user-provided distributions. Event times are managed using spacing variables to enforce chronological order.

For complex parameters, the resize parameters for divergence and bottlenecks are computed as ratios based on the type of event. For divergence events, the first ancestral resize is computed as the ratio of the total ancestral population size (N_ANCALL) to the population-specific ancestral size (N_ANCxy); subsequent divergence resizes are computed as the ratio of the ancestral size of the source-sink pair (N_ANCxy) to the descendant population size (N_POPy). For bottleneck events, the onset resize (RESBOT) is computed as the ratio of the bottleneck size (N_BOTx) to the current population size (N_CURx), and the recovery resize (RESBOTEND) is computed as the ratio of the pre-bottleneck ancestral size (N_ANCx) to the bottleneck size (N_BOTx).

Complex event times are constructed by summing spacing intervals with earlier events to guarantee temporal ordering without constraining absolute values. The first event, whether a divergence, admixture, or bottleneck start event, is assigned directly as a simple parameter drawn from the time distribution. Each subsequent event is defined as a complex parameter by adding a hidden spacing interval to the preceding event: T_2 = T_1_2 + T_1, T_3 = T_2_3 + T_2, and so on, where each spacing parameter T_i_(i+1) is sampled independently from the user-specified spacing range, never exceeding the user-specified maximum time allotted between events. This ensures that all events are strictly ordered in time while allowing the absolute timing of each event to vary freely across the prior.

#### File Naming and Output

Each model is saved to its own subdirectory named {PREFIX}_random_model_N. The .*tpl* and .*est* files are named {PREFIX}.*tpl* and {PREFIX}.*est*.

#### Constraints and Validity

*CoalMiner* enforces temporal and topological consistency during model construction rather than through post-hoc validation. Specifically: (1) events are placed in chronological order by the ordering function during generation; (2) extinct populations are not reused as sources following merger events; and (3) migration matrices are constructed with event-aware connectivity, zeroing out entries for populations that have coalesced at each successive divergence event.

### Language and Performance

*CoalMiner* is written in Python v3.12.2. *CoalMiner* depends on the PyYAML package (v0.2.5). *CoalMiner* is single-threaded and can be run locally or via a high-performance computing environment.

We assessed the runtime of *CoalMiner* as a function of the number of randomly generated models using a three-population configuration. Benchmarks were conducted on a 2023 MacBook Pro equipped with an Apple M2 Pro chip and 16 GB RAM (macOS Sequoia 15.6.1). *CoalMiner* was executed using Python 3 without parallelization.

### Input & Configuration

*CoalMiner* accepts a single YAML configuration file. Required parameters that the user specifies include: INPUT_PREFIX: filename prefixes; NUM_POPS: number of sampled populations; SAMPLE_SIZE: list of haploid sample sizes per population; NUM_RANDOM_MODELS: number of random models to generate (default: 100); prior distributions for mutation rate, effective population sizes, migration rates, and event timing; (optional) max_time_between_events: to constrain event spacing. Observed SFS files must be formatted for *fastsimcoal2* (e.g., *_joint_DAFpop1_0.obs, *_joint_MAFpop1_0.obs) and compatible with the number of populations.

### Output Files

*CoalMiner* generates random .*est* and .*tpl* files and saves them in directories titled {prefix}_random_model_1, {prefix}_random_model_2, etc., in each output directory. It also copies the provided SFS files into the respective model directories. It does not automatically run an entire *fastsimcoal2* analysis, but rather exclusively generates X number of topologies for the user to then test with *fastsimcoal2*.

### Simulation and Testing Framework

To benchmark *CoalMiner*’s ability to recover evolutionary plausible topologies, we used two models: an example simulated model provided with the *fastsimcoal2* package (2PopDivMigr20Mb) and an empirical *Humulus lupulus L.* (hop) dataset (McElwee-Adame et al., 2025). All *fastsimcoal2* analyses were performed with fsc28 (64-bit Linux version; Excoffier et al., 2021).

#### 2PopDivMigr20Mb Data Generation

The 2PopDivMigr20Mb dataset is a two-population divergence-with-migration scenario included as an example dataset with *fastsimcoal2*. This dataset consists of a joint derived allele frequency spectrum computed from 5 haploid samples per population across a 20 Mb locus, and served as a controlled test case for model selection performance. We pulled values from the provided .*est* file as input for *CoalMiner’s* prior distribution ranges. Using those values, along with the provided derived SFS files, we generated 100 random demographic topologies with *CoalMiner*. The parameters used are provided in Supplementary Table 2. Using the .*tpl* and .*est* files from *CoalMiner*, as well as the SFS *.obs* files, we ran 100 separate instances of *fastsimcoal2* for each of the 100 randomly generated models using the following settings (following the recommended settings in Excoffier et al., 2021): *./fsc28 -t hops.tpl -e hops.est -d -0 -C 10 -n 100000 -L 40 -s 0 -M -c 4*, resulting in up to 1,000,000,000 coalescent simulations across all runs.

#### Hop Data Generation

The hop SNP dataset tested here was generated and described in McElwee-Adame et al. (2025). Briefly, genomic DNA was extracted from 163 hop (*Humulus lupulus* L.) accessions and subjected to genotyping by sequencing (GBS) using the ApeKI enzyme, barcoded, and sequenced on an Illumina HiSeq 3000. Reference-free variant calling was performed using the TASSEL 3 GBS pipeline, yielding an initial dataset of 143,309 SNPs. The dataset was subsequently filtered to retain only biallelic sites, remove known triploid cultivars, and exclude sites with a minor allele frequency below 0.05, resulting in a final filtered dataset of 27,163 biallelic SNPs across 98 cultivars spanning four population groups (Central European, English, Noble, and American). Derived allele frequency spectra (DAF) were generated from this filtered VCF using PPP (Webb et al., 2021).

We used these SFS and known domestication history of hops—making sure to include variation in migration rates, population size, admixture rates, and divergence times—to generate 1000 random models with *CoalMiner*. The exact parameters used can be found in Supplementary Table 3. Using the .*tpl* and .*est* files from *CoalMiner*, as well as the SFS *.obs* files, we ran *fastsimcoal2* 1000 times for each of the 1000 randomly generated models using the following settings: *./fsc28 -t hops.tpl -e hops.est -d -0 -C 10 -n 1000 -L 40 -s 0 -M -c 4*, resulting in up to 1,000,000,000 coalescent simulations across all runs.

#### Fastsimcoal2 Estimation

For each dataset, we identified the top-performing models based on the median smallest difference between the MaxEstLhood and MaxObsLhood values across all runs for the model, according to the instructions provided in the *fastsimcoal2* tutorial. We calculated AIC for each model using the formula: *AIC* = 2*k* – 2*ln*(*L*) where *k* is the number of free parameters (as defined in the .*est*) and *L* is the maximum likelihood of the model.

To assess the stability and reliability of the likelihood for each candidate model, we resampled the likelihood distribution of the best-fit parameter estimates using *fastsimcoal2*. For each model, the maximum likelihood parameter estimates (stored in the *_maxL.par* file from the initial optimization runs) were used as fixed input parameters, and *fastsimcoal2* was run 100 times with 100,000 simulated genealogies per iteration (-n 100000) to obtain a distribution of log-likelihoods evaluated at those parameter values. This approach characterizes the stochastic variance in the SFS-based likelihood approximation that arises from the coalescent simulations underlying *fastsimcoal2*’s expectation computation. Models were ranked and compared using the mean and maximum log-likelihood across the 100 iterations. We exclusively used those top-performing models for downstream analyses, specifically the highest-performing runs for each model.

### Confidence Intervals via Parametric Bootstrap

Upon successful run completion, *fastsimcoal2* creates a *_maxL.par* file per run instance, which is created from the maximum likelihood parameters that were estimated during the run. Following the procedures described in the *fastsimcoal2* manual, we used the *_maxL.par* file from each model’s best run to perform parametric bootstrapping. We modified the file to replace “FREQ 1” with “DNA 100” and set the number of loci to “200000 0,” representing 200,000 non-recombining segments of 100 base pairs. Using *fastsimcoal2*, we then simulated 100 pseudo-observed SFS replicates per model: *./fsc28 -i prefix_boot.par -n 100 -j -d -s 0 -x -I -q -c 4*.

Each bootstrap replicate was then re-analyzed using the same topology and estimation settings, now with the pseudo-observed SFS, with 100 *fastsimcoal2* runs per replicate, with the following settings: *./fsc28 -t prefix_boot.tpl -e prefix_boot.est -d -0 -C 10 -n 1000 -L 40 -s 0 -M -c 4*. We then extracted the best run from each replicate by comparing the maximum observed likelihood and the maximum estimated likelihood and compiled parameter distributions to derive 95% confidence intervals.

### Assumptions and Model Constraints

All demographic inferences were performed under the composite likelihood framework implemented in *fastsimcoal2*, which summarizes genomic data as the SFS and estimates the likelihood of demographic parameters by treating each single nucleotide polymorphism (SNP) as an independent observation under a neutral coalescent model, thereby ignoring linkage among sites.

Demographic models generated by *CoalMiner* were constrained to ensure logical consistency of evolutionary histories. Specifically, populations were not permitted to reappear following merger events, and historical events were enforced to occur in a temporally consistent order. Migration was modeled using event-aware connectivity matrices, restricting gene flow to biologically plausible population configurations.

Parameter estimation and model evaluation, therefore, reflect these assumptions and constraints, and inference accuracy is contingent on the extent to which the simulated and empirical data conform to them.

### Software and Data Availability

All scripts, detailed instructions for using *CoalMiner,* and examples for downstream analyses, SFS files, and other relevant data are publicly available on GitHub: https://github.com/raywray/CoalMiner and https://github.com/raywray/CoalMiner_Paper_Validation.

## Results

### Performance

Runtime increased approximately with the number of models generated, ranging from 3.5 seconds for 10 models to 476 seconds for 5000 models. Runtimes were measured using wall-clock time from a shell script invoking *CoalMiner* with a fixed YAML configuration file. A full breakdown of runtimes is provided in Supplementary Table 1, and the scaling relationship is shown in Supplementary Figure 1.

### *CoalMiner* generates diverse demographic topologies

To evaluate the diversity of demographic models generated by *CoalMiner*, we examined the topological composition of 100 randomly generated models for a two-population system (2PopDivMigr20Mb). *CoalMiner* produced substantial variation in both the presence and absence of demographic features and in the number and type of historical events per model (Figure 2). Ghost populations were present in 59% of models, while admixture events occurred in 51% of models. Migration between demes was included in 51% of models, with 23 of those featuring migration restricted to a single time period (one non-zero migration matrix) and 28 featuring migration across two distinct periods.

**Figure 2:**
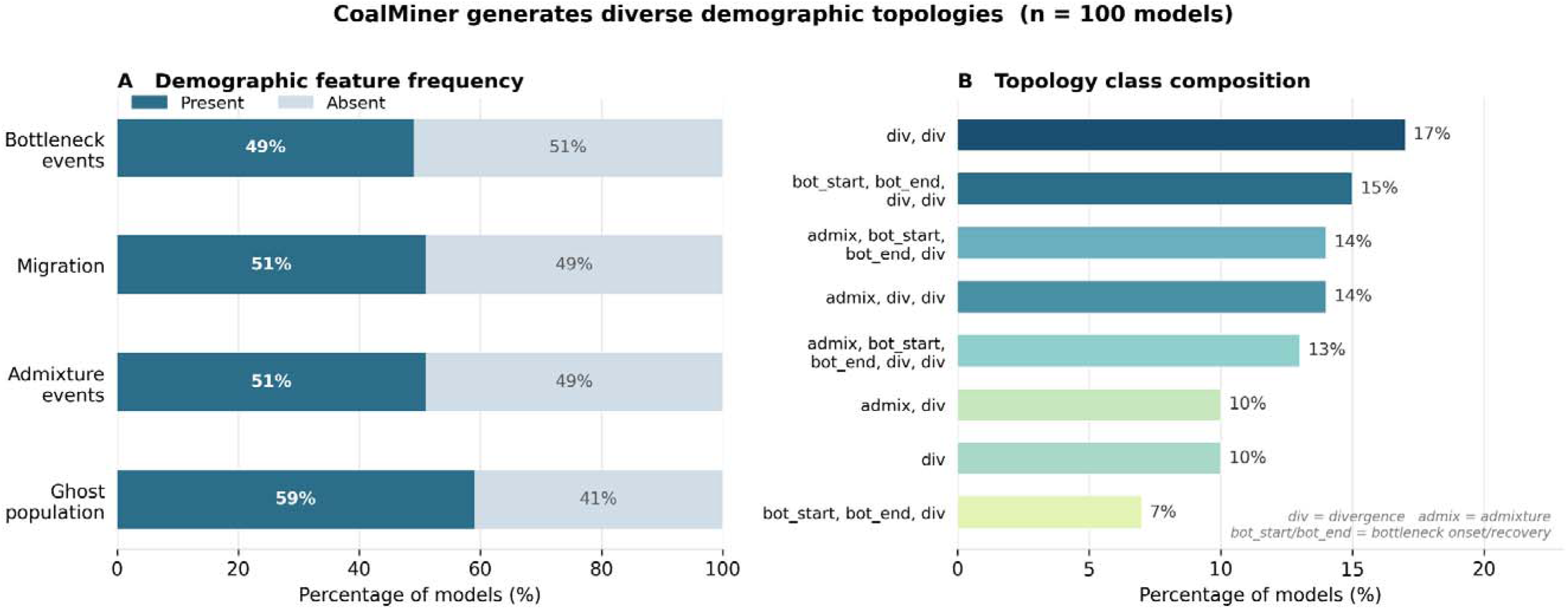
*CoalMiner* generates diverse demographic topologies across 100 randomly generated models for the two-population 2PopDivMigr20Mb benchmark dataset. (A) Demographic feature frequency, showing the proportion of models that contain each event type. (B) Topology class composition showing the distribution of models by their unique combination of event types. Abbreviations: div = divergence event; admix = admixture event; bot_start/bot_end = bottleneck onset and recovery event.

Bottlenecks were present in 49% of models. In terms of event complexity, models ranged from a single historical event to five events, with the plurality (29%) containing four events. Across all 100 models, *CoalMiner* generated 159 divergence events, 51 admixture events, and 49 bottleneck onset and recovery events. Eight distinct topology classes emerged based on event composition, the most common being simple two-divergence models (17%), followed by models combining divergence with bottlenecks (15%) and divergence with admixture (14%). No single topology dominated the ensemble, and all 16 possible combinations of ghost population, admixture, migration, and bottleneck presence/absence were represented, demonstrating that *CoalMiner* explores a broad and demographically plausible model space without systematic bias toward any particular topology.

For the hops analysis, we only retained the top ten performing models for each downstream analysis.

### *CoalMiner* recovers high-likelihood models for the 2PopDivMigr20Mb benchmark

To evaluate *CoalMiner’s* ability to identify well-fitting demographic models from a large candidate space, we applied it to the 2PopDivMigr20Mb benchmark dataset. From 100 randomly generated *CoalMiner* models, we evaluated fit using composite log-likelihood and AIC distributions computed across 100 *fastsimcoal2* likelihood evaluation replicates per model, with the number of free parameters (k) estimated from each model’s .*est* file.

Interestingly, when AIC was computed accounting for model complexity, the ranking of models changed substantially relative to likelihood alone: models with the highest likelihoods (random_model_23, k=18; random_model_17, k=19) ranked worst by AIC, as their additional parameters provided no meaningful improvement in fit. The two best-supported models by AIC were random_model_52 and random_model_87, both with k=9, median AIC values of 20,275.59 and 20,275.66, respectively, and AIC distributions clearly separated from all other models (Figure 3; Table 1). The third-ranked model, random_model_79 (k=10, median AIC 20,277.54), included an admixture event and ranked ∼2 AIC units above the top two. Models with k≥14 were separated from the top group by ≥9 AIC units, providing strong support for the simpler topologies. Convergence of the likelihood optimizer was generally good across the top models, with AIC distribution ranges of less than 2 units (random_model_52 range: 1.85; random_model_87 range: 1.75), indicating stable and reproducible optimization across replicates.

**Figure 3:**
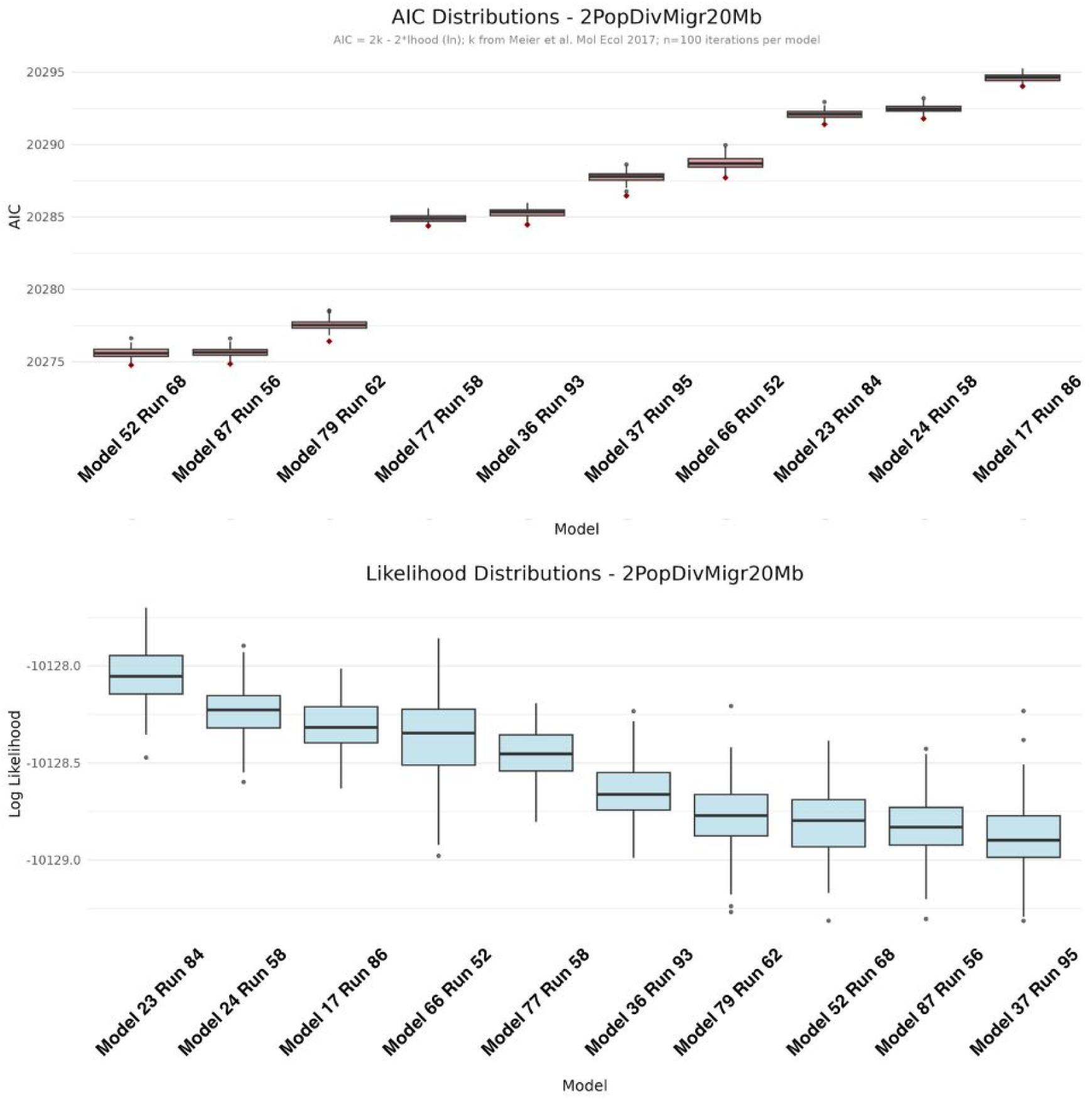
AIC and log-likelihood distributions across the top-performing *CoalMiner* models for the 2PopDivMigr20Mb benchmark dataset (n = 100 likelihood evaluation iterations per model). (A) AIC distributions ranked from lowest (best fit) to highest. (B) Likelihood distributions for the same models ranked from highest to lowest, shown for comparison. Red points indicate outliers.

**Table 1:** Model comparison summary for the top-performing *CoalMiner* models evaluated on the 2PopDivMigr20Mb benchmark dataset, ranked by median AIC across 100 likelihood evaluation replicates.

| Model | k | Median_lhood | Median_AIC | Min_AIC | AIC_range | Bootstrap_TDIV_median | Bootstrap_TDIV_95CI_lower | Bootstrap_TDIV_95CI_upper | Notes |
| --- | --- | --- | --- | --- | --- | --- | --- | --- | --- |
| random_model_52 | 9 | -10128.8 | 20275.59 | 20274.77 | 1.85 | 54527 | 2471 | 118257 | Top model; core div+migration topology |
| random_model_87 | 9 | -10128.83 | 20275.66 | 20274.85 | 1.75 | 61135 | 3940 | 130883 | Top model; core div+migration topology; time-varying migration |
| random_model_79 | 10 | -10128.77 | 20277.54 | 20276.41 | 2.12 | 93029 | 6303 | 166802 | 3rd; includes admixture event (T_ADMIX01 median 58512; 95CI 3139-126602) |
| random_model_77 | 14 | -10128.45 | 20284.91 | 20284.38 | 1.22 | 135655 | 37601 | 223414 | Best k=14 model; includes bottleneck; comparison case only |
| random_model_23 | 18 | -10128.05 | 20292.11 | 20291.4 | 1.54 | — | — | — | Highest lhood but worst AIC; overparameterized |
| random_model_17 | 19 | -10128.32 | 20294.63 | 20294.03 | 1.23 | — | — | — | Highest k; worst AIC; overparameterized |

To characterize parameter uncertainty in the best-supported models, we performed parametric bootstrap analyses with 100 replicates on random_model_52, random_model_87, random_model_79, and random_model_77 (the best k=14 model, included for comparison). All four models shared a core topology of two sampled populations connected by bidirectional migration and a single ancestral divergence event, with random_model_79 additionally including an admixture event and random_model_77 including a population bottleneck. Across all models, divergence time (T_DIV) was the most consistently estimable parameter (Figure 4): random_model_52 recovered a median T_DIV of 54,527 generations (95% CI: 2,471–118,257), and random_model_87 a median of 61,135 generations (95% CI: 3,940–130,883), with both 95% CIs spanning roughly one order of magnitude. In random_model_79, the divergence time estimate was deeper (median T_DIV: 93,029 generations; 95% CI: 6,303–166,802) and was accompanied by an admixture time estimate of 58,512 generations (95% CI: 3,139–126,602), suggesting a plausible admixture-after-divergence history, though with substantial uncertainty. Population size parameters (N_POP0, N_POP1) showed considerably wider uncertainty, with 95% CI widths spanning 2–3 orders of magnitude in all models, consistent with the well-documented difficulty of estimating absolute effective sizes from small-sample SFS data. Migration rate parameters were effectively unidentifiable across all models: median estimates for both MIG01 and MIG10 were near zero (∼10) with 95% CIs spanning many orders of magnitude, indicating that the SFS of this benchmark dataset carries insufficient signal to distinguish migration from zero. The relative ancestry parameter (RELANC01), which encodes the ancestral population size as a proportion of the descendant population, was similarly poorly constrained. Taken together, these results indicate that *CoalMiner* successfully identifies parsimonious, well-fitting demographic topologies via AIC-based model selection, while the bootstrap analyses reveal that divergence time is the most recoverable parameter in this two-population system, given the available data.

**Figure 4:**
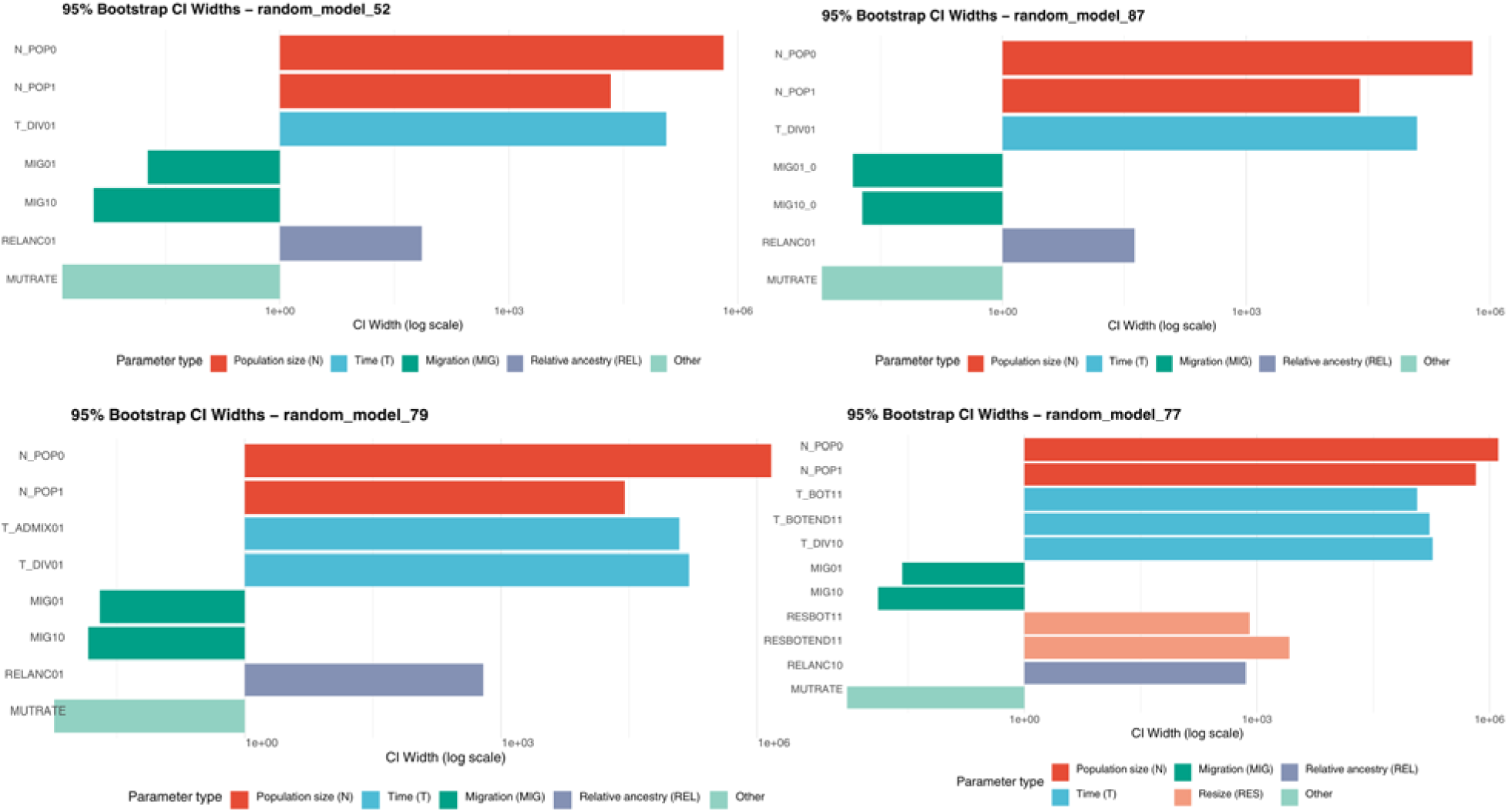
95% bootstrap confidence interval widths for estimated demographic parameters across the four candidate models selected from the 2PopDivMigr20Mb benchmark analysis (random_model_52, random_model_87, random_model_79, and random_model_77), based on 100 parametric bootstrap replicates per model. Bar length reflects the width of the 95% CI on a log scale, where longer bars indicate greater parameter uncertainty. Parameters are colored by type: population size (N, red), divergence and event times (T, blue), migration rates (MIG, dark green), relative ancestry (REL, purple), bottleneck resize parameters (RES, salmon), and other parameters including mutation rate (light green). random_model_77 includes additional parameters reflecting its more complex topology, which incorporates a bottleneck event.

### *CoalMiner* recovers biologically meaningful demographic models from empirical hop data

*CoalMiner* generated demographic models for domesticated hop cultivars that included multiple divergence, bottleneck, admixture, and migration events consistent with previously inferred breeding histories of cultivated hops. A total of 1000 randomly generated demographic models were evaluated using *fastsimcoal2* likelihood optimization and ranked using likelihood differences and AIC estimates. The highest-performing model recovered four major cultivar groups corresponding to Central European, Noble, English, and American ancestries and included multiple historical migration events between lineages (consistent with McElwee-Adame et al. (2025); Figure 5). Estimated divergence events included an early split between Central European and American cultivar groups approximately 2799 years before present (95% CI: 2724–2799 ybp), followed by divergence of Noble cultivars approximately 2337 ybp (95% CI: 2353–2430 ybp), and a more recent divergence associated with English cultivars approximately 624 ybp (95% CI: 630–648 ybp).

**Figure 5:**
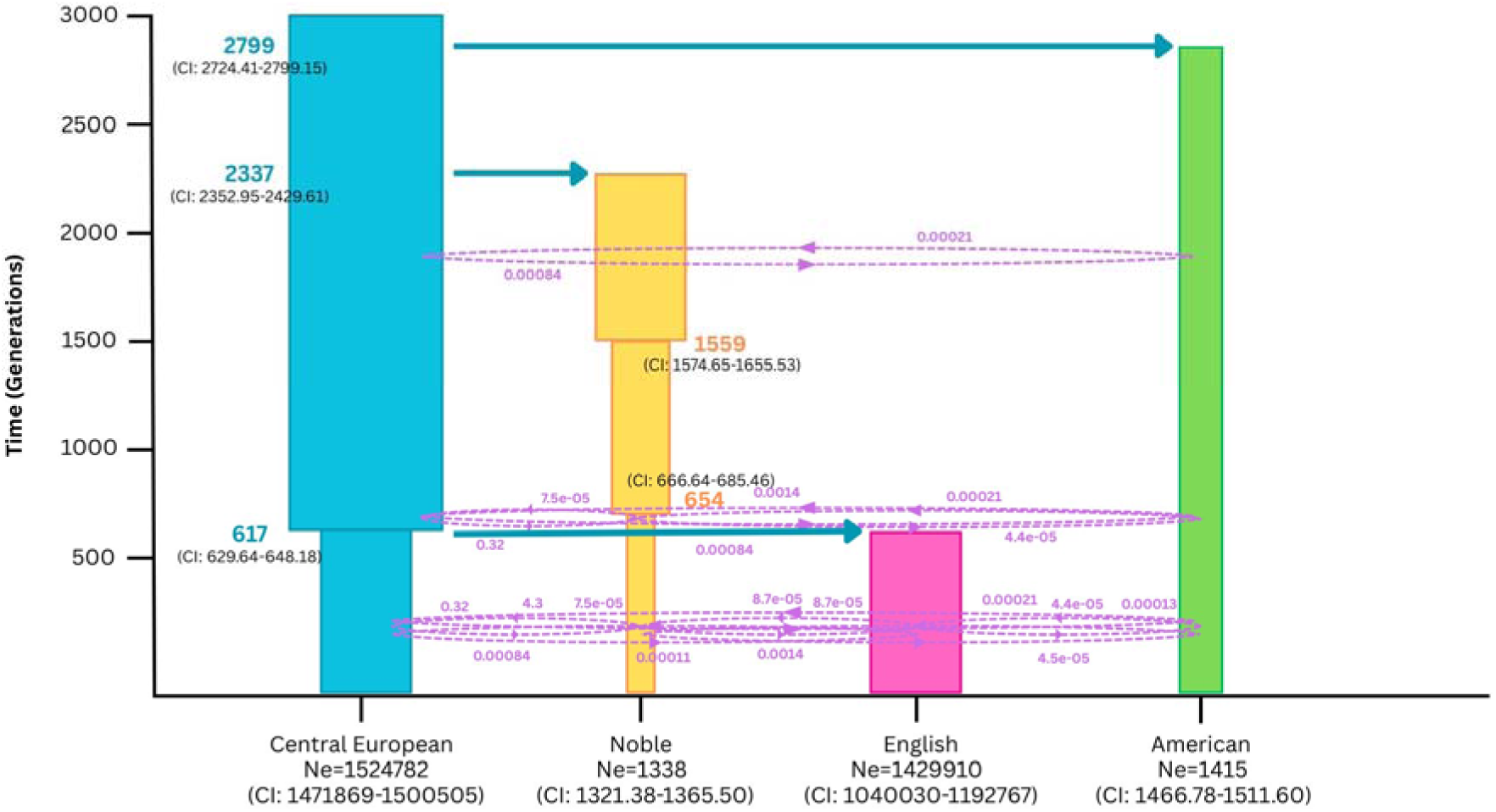
Most likely model of evolutionary history for the four predominant provenances determined by *CoalMiner* and *fastsimcoal2* analyses. Group 1 (pink) = Central European Ancestry, Group 2 (green) = Noble Ancestry, Group 3 (orange) = English Ancestry, Group 4 (light blue) = American Ancestry. Estimates of divergence times are shown in blue font; estimates of bottleneck times are shown in yellow. The most likely model includes several significant migration events, here shown in purple dotted lines/arrows.

The inferred demographic history additionally recovered strong bottleneck signatures in American, Central European, and Noble cultivar groups, with substantially reduced contemporary effective population sizes relative to ancestral populations. Parametric bootstrap analyses recovered stable confidence intervals across major demographic parameters. Together, these results demonstrate that *CoalMiner* can recover biologically coherent demographic histories from empirical population genomic data.

Additionally, the estimated demographic history accurately captures known historical records in the species (McElwee-Adame et al., 2025).

## Discussion

*CoalMiner* is a useful tool designed to help overcome the limitations of manual user-provided topologies to demographic modeling software, specifically *fastsimcoal2*. Not only does the tool aid in the sheer number of models to test, but it also provides a diverse array of plausible models. Additionally, we anticipate that it will help researchers simultaneously save time in learning about the input files for such software, while also allowing for the exploration of possible model space.

Through our benchmarking and empirical analyses, we show that *CoalMiner* is extremely useful when paired with any prior model knowledge. In our two-population model, *CoalMiner* was able to accurately recover plausible demographic parameters and events. Further, the empirical analysis of hops allowed for stronger support of the Noble lineage as distinct and unique.

*CoalMiner’s* strengths lie in its user-friendliness, run time, and simplistic algorithm (it isn’t over-engineered). However, its simple decision tree-based pipeline can also be a flaw. Real biology is complicated and complex, so there is an argument that randomized choices to add or not to add events, populations, and migrations can oversimplify results. To address that concern, we advise that *CoalMiner* not be a one-stop shop for demographic analysis, but rather use it as a first stop to identify parameters or events that are highly supported that were perhaps not considered in the first place, and use that to fine-tune analyses.

As an additional note, while SFS-based demographic inference has allowed researchers to evaluate evolutionary histories with much less computational intensity and can be a wonderfully informative summary statistic, we would like to caution end-users of two outstanding issues with inference of evolutionary demographic history using the SFS (or versions of it):

a. Multiple distinct demographic histories can produce nearly identical site frequency spectra, representing a fundamental identifiability limitation of SFS-based inference. This non-identifiability arises from several sources. First, different demographic scenarios (such as deep divergence with continued gene flow versus recent divergence without gene flow) can yield statistically indistinguishable SFS shapes (Lapierre et al., 2017; Marchi et al., 2024). Second, and more broadly, selection at linked sites can generate SFS patterns that closely mimic demographic signals: Johri et al. (2021) demonstrate that background selection creates a skew toward rare alleles that is nearly indistinguishable from population growth, such that neglecting background selection results in false inference of growth, regardless of the true demographic history. Additionally, inbreeding and population size changes both affect the low-frequency entries of the SFS in ways that are frequently confounded (Blischak et al., 2019). Therefore, while *CoalMiner* can help with a broad assessment of the breadth of evolutionary models that capture the observed SFS, any meaningful inferences about the ground truth can only be made with the biology and phylogeographic history of the system firmly in mind.
b. Correspondingly, we have also shown that overparametrized models can very well result in highly likely models, often better than the ‘true’ model. This raises a deeper question about the identifiability of complex evolutionary parameters from the shape of the SFS alone. This problem is compounded by the fact that SFS-based composite likelihood methods treat SNPs as independent observations, which overestimates the true number of independent sites and leads to underpenalization of parameter-rich models under standard model selection criteria such as AIC (Adrion et al., 2020; Johri et al., 2021). As a result, AIC rankings in our framework, and in SFS-based inference generally, should be interpreted as a guide for relative model comparison rather than as definitive model selection. We therefore recommend utilizing a combination of lines of evidence prior to making inferences, especially with larger topologies involving higher-order parameters.

Additionally, we caution users that since *CoalMiner* implements a randomized framework to generate models, some models may result in segmentation faults or memory issues while running *fastsimcoal2*.

Together, however, we suggest that *CoalMiner* is capable of recovering near-optimal models that closely match true demographic scenarios, especially when model complexity is moderate, and the signal in the SFS is strong.

## Acknowledgments

This work was supported by NSF ABI 1564659, NSF CAREER 2147812, and NIH 1R15GM143700-01 to AS. RES was supported via startup monies to AS at San Diego State University, DE-SC0025673 to PI Anand (SFSU) and co-PI Sethuraman, and an SDSU University Graduate Fellowship. All computations were performed on the *mesxuuyan* and *anthill* HPC clusters at San Diego State University that were supported by startup funds to AS, at the UC Riverside HPC, and the SDSC Expanse HPC at UC San Diego. We would like to thank Stephanie Young, Trevor Mugoya, and members of the Sethuraman Lab for their useful discussions on this manuscript.

## Declaration of use of Generative AI

Claude Sonnet 4.6 (Anthropic, accessed 2025–2026) was used to assist with manuscript editing and the generation of analysis scripts. All AI-assisted content and code were reviewed, tested, and verified by the authors, who take full responsibility for the accuracy of all results and interpretations.

