## Supplemental Material for "CoalMiner: a coalescent model generator for *fastsimcoal2*"

### Supplementary Figures and Tables

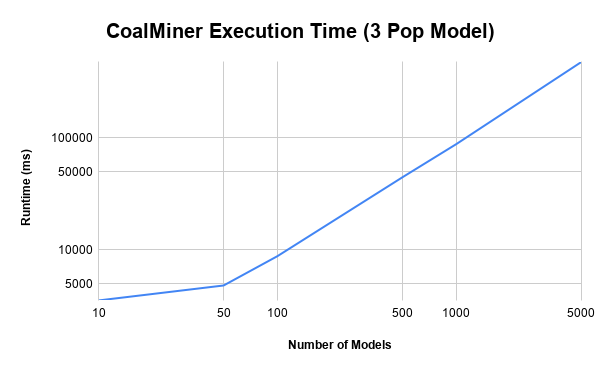

Figure 1: *CoalMiner* runtime as a function of the number of models generated for a three-population configuration, measured in milliseconds on a 2023 MacBook Pro (Apple M2 Pro, 16 GB RAM) without parallelization. Runtime scales approximately linearly with the number of models generated.

| Number of Models | Runtime (ms) |
| --- | --- |
| 10 | 3532 |
| 50 | 4804 |
| 100 | 8782 |
| 500 | 44469 |
| 1000 | 87924 |
| 5000 | 475602 |

Table 1: CoalMiner wall-clock runtimes for generating between 10 and 5,000 demographic models under a three-population configuration.

Table 2: *CoalMiner* prior distributions and fastsimcoal2 run settings used for the 2PopDivMigr20Mb benchmark analysis. Parameter bounds were derived from the values provided in the example .*est* file distributed with *fastsimcoal2*.

| Parameter | Distribution | Minimum | Maximum | Notes |
| --- | --- | --- | --- | --- |
| --- General configuration --- |  |  |  |  |
| Num. populations | — | 2 | 2 | Fixed |
| Sample size (pop. 1) | — | 5 | 5 | Haploid individuals; fixed |
| Sample size (pop. 2) | — | 5 | 5 | Haploid individuals; fixed |
| --- Model parameters --- |  |  |  |  |
| Mutation rate | Uniform | 2.50E-08 | 2.50E-08 | Per site per generation; fixed |
| Effective pop. size (N) | Log-uniform | 10 | 1.00E+07 | Haploid individuals |
| Migration rate | Log-uniform | 1.00E-10 | 1.00E-01 | Per gene copy per generation |
| Divergence time (T) | Uniform | 10 | 1.00E+05 | Generations |
| Max. time between events | — | — | 1.00E+05 | Generations; constrains event spacing |
| --- Run settings --- |  |  |  |  |
| Num. random models | — | — | — | 100 |
| Num. fsc28 runs per model | — | — | — | 100 |
| Coalescent sims per run | — | — | — | 100000 (-n) |
| ECM iterations | — | — | — | 40 (-L) |
| Threads | — | — | — | 4 (-c) |

Table 3: *CoalMiner* prior distributions and *fastsimcoal2* run settings used for the empirical hop analysis. Sample sizes reflect the haploid counts per population extracted from the filtered VCF.

| Parameter | Distribution | Minimum | Maximum | Notes |
| --- | --- | --- | --- | --- |
| --- General configuration --- |  |  |  |  |
| Num. populations | — | 4 | 4 | Fixed; Central European, English, American |
| Sample size (pop. 1) | — | 18 | 18 | Haploid individuals; from VCF |
| Sample size (pop. 2) | — | 19 | 19 | Haploid individuals; from VCF |
| Sample size (pop. 3) | — | 41 | 41 | Haploid individuals; from VCF |
| Sample size (pop. 3) | — | 20 | 20 | Haploid individuals; from VCF |
| --- Model parameters --- |  |  |  |  |
| Mutation rate | Uniform | 6.10E-09 | 6.10E-09 | Per site per generation; fixed. Based on Arabidopsis thaliana rate (Ossowski et al. 2010) |
| Effective pop. size (N) | Uniform | 10 | 2.00E+06 | Haploid individuals |
| Migration rate | Log-uniform | 1.00E-05 | 5 | Per gene copy per generation |
| Divergence time (T) | Uniform | 1 | 5000 | Generations |
| --- Run settings --- |  |  |  |  |
| Num. random models | — | — | — | 1000 |
| Num. fsc28 runs per model | — | — | — | 1000 |
| Coalescent sims per run | — | — | — | 1000 (-n) |
| ECM iterations | — | — | — | 40 (-L) |
| Threads | — | — | — | 4 (-c) |
